# A novel function of *neuroligin 3*: Beyond sociality, Regulating lifespan in *Drosophila*

**DOI:** 10.64898/2026.10.27.762327

**Authors:** Judy M. Kurbaj, Reza Ataei, Anne F. Simon

## Abstract

Neuroligins are conserved postsynaptic adhesion proteins involved in synaptic function and behaviour. In *Drosophila melanogaster*, Neuroligin 3 (Nlg3) regulates social behaviour, sleep and synaptic architecture. We now report, through analysis of publicly available transcriptomics data, confirmed with RT-qPCR data, that loss-of-function of the *nlg3* gene downregulates the expression of longevity-associated genes *foxo*, *hsp7*0s, and *hsp68*. Graded genetic manipulation of the *Drosophila melanogaster neuroligin 3* (*nlg3*) gene indeed modulates both social spacing and lifespan. Specifically, reduced *nlg3* expression (via heterozygous deletion, or transposable-element insertion) extends lifespan in both sexes, while overexpression or complete deletion shortens it. In contrast, overexpression or complete loss of *nlg3* leads the flies to be further apart (increased social distance, or social space), as does RNAi knockdown, while heterozygous reduction of *nlg3* expression either lowers or does not alter the flies’ social space. This work identifies a previously unrecognized link between a conserved synaptic gene, social behaviour, and organismal longevity.

## Introduction

Social behaviour is a widespread feature of animal life, shaping survival and reproductive success through interactions among conspecifics [1,2]. Across evolutionary lineages, these interactions have given rise to diverse forms of cooperation and coordination, ranging from the division of labour in eusocial insects to enduring social bonds in mammals and collective movement in schooling fish [3–5]. Such behaviours can influence resource acquisition, predator avoidance, and reproductive opportunities, thereby contributing to individual fitness [6,7]. Beyond these immediate ecological and reproductive consequences, social organization has also been associated with variation in lifespan [8,9]. In group-living animals, social relationships can improve access to resources, reduce predation risk, support reproduction, and buffer physiological stress, while other social interactions can impose energetic or reproductive costs [10–12]. In *Drosophila melanogaster*, females with higher early-life mating rates tend to have shorter lifespans [13], consistent with a trade-off between reproductive investment and later survival [14]. A comparative analysis of approximately 1,000 mammalian species found that group-living species generally exhibited greater longevity than solitary species [15]. These observations suggest that social organization and longevity may be evolutionarily associated, raising questions about the biological mechanisms underlying this relationship.

Aging is characterized by a progressive decline in physiological function and reproductive capacity, ultimately increasing the risk of mortality [16]. Although lifespan is strongly influenced by conserved genetic pathways that regulate metabolism, stress resistance, and cellular maintenance, behaviour can also shape survival. In *Drosophila melanogaster*, social and reproductive experiences have measurable consequences for lifespan, suggesting that behavioural state and aging are not two independent processes. More broadly, social and sensory experiences can alter survival in flies, raising the possibility that neural pathways controlling behaviour may also influence physiological processes associated with aging [17–19]. Social spacing provides a useful behaviour in which to examine this relationship. Social spacing refers to the preferred distance individuals maintain from conspecifics and depends on the detection and integration of attractive and repulsive social cues [20]. In *Drosophila*, this behaviour is reproducible, genetically tractable, and influenced by sex, genetic background, dopamine signalling, and specific neural circuits [21–23]. Although studies have shown that social experience and neural signalling can influence lifespan, the genetic mechanisms linking baseline social behaviour with longevity remain poorly understood.

Neuroligins are conserved postsynaptic adhesion proteins that interact with presynaptic Neurexins to regulate synapse organization and function [24,25]. Despite evolutionary divergence between vertebrate and invertebrate Neuroligins, important structural and functional properties are conserved across species [26]. In humans, variants in *NLGN3* and *4* have been associated with autism spectrum disorder and altered social communication [27], while manipulation of *neuroligin* genes (*nlgs*) in *Drosophila* affects social spacing, aggression, sociability, and sleep [28–31]. Previous work has specifically identified one of the 4 Drosophila *nlgs* genes, Drosophila *neuroligin 3* (*nlg3*) as a regulator of adult Drosophila social neural substrates. Antibodies and reporter of expression constructs localized protein and gene expression to specific brain regions: the mushroom-body calyx, optic lobes, and protocerebral bridge [32]. RNAi knockdown implicated cholinergic, dopaminergic, and sexually dimorphic *fruitless*-positive neurons in a sex-specific manner [32]. The Nlg3 protein is required for normal neuromuscular-junction architecture and synaptic-vesicle endocytosis [33]. Nlg3 undergoes constitutive, TACE-dependent proteolytic maturation required for normal locomotion [34], and an Nlg3 C-terminal fragment can enter the nucleus in cultured cells and regulate innate-immune genes [35]. And a recent report further shows fly *nlg3* gene deletion disrupts sleep and synaptic architecture, So *nlg3* loss-of-function not only alters social behaviour, but also disturbs sleep architecture and immune response, two traits mechanistically linked to aging [36–38]. More specifically, Xie et al. (2023) [35] report the role of *nlg3* in controlling the immune response: that *nlg3* null flies display an activation of the innate immune responses, and an increased expression of the innate immune response genes. Their differential expression analysis identified 472 significantly altered genes, of which 312 were upregulated and 160 were downregulated in *nlg3* knockout flies compared with wild-type controls. Gene Ontology analysis showed that the upregulated genes were strongly enriched for innate immune response and defense-related pathways, particularly responses to bacterial infection. Consistent with the RNA-seq results, several immune-related genes associated with the IMD and Toll pathways, including *Dpt, Mtk, Drs, Im1, AttA, AttC, CecA, CecB,* and *CecC*, were subsequently confirmed to be upregulated by RT-qPCR. These findings indicate that loss of *nlg3* produces broad transcriptional changes in the adult fly head, with a particularly strong effect on genes involved in innate immune regulation.

Here, we reanalyzed those transcriptomics data, made publicly available by Xie et al. (2023) [35] and found enrichment in longevity-related genetic pathways in their *nlg3* loss-of-function flies. We confirmed those results through RT-qPCR on our own deficiency flies, as well as on an allelic series of graded manipulation of *nlg3* expression: *nlg3* overexpression, RNAi-mediated knockdown, and heterozygous deficiency. We further phenotyped this allelic series and found that increasing *nlg3* expression increased social spacing and reduced survival in both sexes, whereas reducing *nlg3* expression extended lifespan, with the behavioural effects differing between heterozygous deficiency and RNAi-mediated knockdown. Together with changes in *foxo*, *hsp68*, and *hsp70*, these findings support that *nlg3* acts as a dosage-sensitive regulator of social behaviour and lifespan in male and female *Drosophila* and identify proteostasis-related pathways as a candidate molecular connection between synaptic function and organismal aging.

## Materials and Methods

### Fly lines

Full genotypes and stock information for all fly lines used for RT-qPCR analysis, longevity and behavioural experiments, are provided in **Table 1**. To control for genetic background effects, all lines used for RT-qPCR, longevity and behavioural analysis were outcrossed 5 times to the laboratory control strain Canton-S (Cs) before experimentation. The genotype of the fly lines that generated the transcriptomics data is from Xie et al. (2022) [35].

**Table 1.** Parental Drosophila stocks used to generate experimental fly lines. Stock names were simplified for use throughout the text. BL: Bloomington Stock Center number.

| Stock name | Stock number | Genotype | Chromosomal location | Type |
|---|---|---|---|---|
| **Cs** | In-house | Control genotype used for all outcrosses | - | Control |
| **UAS-DNA<sup>nlg3</sup>** | In-house [28] | w<sup>1118</sup>; UAS- cDNA<sup>nlg3</sup> Outcrossed 5 times in Cs | 1;2 | UAS effector Outcrossed 5 times in Cs |
| **UAS-RNAi<sup>nlg3</sup>** | In-house [23] | w+; UAS-nlg3 RNAi Outcrossed 5 times in Cs | 1;2 | UAS effector Outcrossed 5 times in Cs |
| ***nlg3-Gal4*** | BL 76134 | w<sup>1118</sup>; Mi{Trojan-GAL4.2}Nlg3[MI00445-TG4.2]/TM3 | 1;;3 | Trojan-Gal4-driver Outcrossed 5 times in Cs |
| ***nlg3<sup>def</sup>*** | In-house [28][39] | w<sup>1118</sup>;; Df(3R)nlg3 IID-24 (w+) Outcrossed 5 times in Cs | 1;;3 | Deletion of *nlg3* Outcrossed 5 times in Cs |
| ***nlg3<sup>KO88</sup>*** | Xie et al. (2023) [35]<br>Xing et al. (2014) [40] |  | 1;;3 | Knockout of *nlg3* |
| ***nlg3<sup>KO127</sup>*** |  |  | 1;;3 | Knockout of *nlg3* |
| ***w<sup>1118</sup>*** |  |  | 1 | Control used for transcriptomics |

### Fly husbandry

All lines were maintained in mixed-sex bottles containing standard media prepared in-house (same recipe as Fisherbrand™ Jazz-Mix™ Drosophila Food). Flies were reared in the insect module of the Biotron Facility at University of Western Ontario under controlled environmental conditions of 25°C, 50% relative humidity, and a 12:12 light-dark cycle.

### Re-analyzing RNAseq publicly available data

The Gene Expression Omnibus (GEO) is a publicly accessible repository for gene expression and functional genomics data [41]. Publicly available RNA-sequencing data generated by Xie et al. (2023) [35] were obtained from GEO under accession GSE219064. The dataset consists of six adult *Drosophila melanogaster* head RNA-seq samples representing two genotypes: three biological replicates of wild-type (*w*1118) controls (GSM6766965 - GSM6766967) and three biological replicates of *nlg3* knockout flies (*nlg3*KO88/*nlg3*KO127; GSM6766968 - GSM6766970). In the original study, total RNA was extracted from pooled adult fly heads using TRIzol, and high-throughput sequencing was performed using the Illumina HiSeq 2000 platform [35].

We imported raw sequencing data into the Galaxy platform for reanalysis. The quality of the raw sequencing reads was first assessed using FastQC [42]. Reads were subsequently processed using Trimmomatic [43] to remove universal adapter sequences and filter low-quality and short reads, using a minimum read length of 30 bp and a quality threshold of 20. The filtered reads were aligned to the *D. melanogaster* Dm6 reference genome [44] using HISAT2 [45], and gene-level read counts were generated using featureCounts [46] using dmel-all-r6.60.gtf.gz annotation file (compressed Gene Transfer Format (GTF) genomic annotation file for Drosophila melanogaster, released by FlyBase [47]. Differential gene expression between *nlg3* knockout and wild-type samples was subsequently analyzed using DESeq2 [48]. Genes with an absolute log₂ fold change (|log₂FC|) ≥ 1.3 and a *p*-value < 0.05 were considered differentially expressed. KEGG pathway enrichment analysis was performed on the downregulated gene set using FlyEnrichr [49], the *Drosophila melanogaster*-specific implementation of Enrichr. The KEGG pathway library was used to identify enriched pathways [49].

### Reverse-transcription quantitative polymerase chain reaction (RT-qPCR)

For each sample, heads were collected from approximately 20 flies aged 3–5 days. Control and treatment flies matched the genotypes and ages used in the longevity assays. Total RNA was isolated with TRIzol reagent (Invitrogen, #15596026) following the manufacturer’s protocol and treated with TURBO DNase (Invitrogen AM2238) to remove genomic DNA. cDNA was prepared using the iScript cDNA Synthesis Kit (#1708841). RT-qPCR was carried out with PowerUp SYBR Green Master Mix (Thermo Fisher Scientific) on a Bio-Rad CFX Connect Real-Time PCR Detection System (Bio-Rad Laboratories, Hercules, CA, USA) to determine the relative gene expression levels of *hsps* and *foxo* transcripts. Primer sequences are provided in

Relative expression levels of transcripts was determined by the ΔΔCt method , using α*Tubulin 84* (*αTub84B*) as the reference gene, for 3-4 technical and 3 to 4 biological replicates.

### Handling prior to behaviour and longevity assays

All flies were maintained in mixed-sex social groups. Newly eclosed flies were collected from stock bottles and transferred to fresh food bottles for aging for 3-4 days (most of the flies are no longer virgin at that age [50]). All behavioural assays were conducted in the same room under controlled temperature and humidity (25°C and 50%humidity), as well as standardized lighting conditions between 12:00 PM and 4:00 PM (Zeitgeber time 4–8) to minimize variation associated with circadian rhythms.

### Social space behaviour assays

The social space assay was performed as described by Simon et al (2012) and McNeil et 2015 [20,22]. Twenty-four hours prior to the assay, flies were sexed under cold anesthesia and separated into groups of 12 to 17 in vials with food for each treatment tested. Two hours before the assay, each treatment was transferred into new vials and acclimated to standard conditions. At the time of testing, flies were loaded into the triangular chamber and knocked to the bottom of the chamber to induce a startle response that quickly forces the flies into a group at the top of the chamber. Flies were then given 20 to 40 min to settle at their preferred social space. Each treatment was loaded into separate social space chambers and photos were taken as soon as flies had settled. For more details, see the video associated to the protocol description, in McNeil et al. (2015) [22]. Images were analyzed in ImageJ using plugins developed by Yost et al. (2020) [51]. Social space was quantified as the mean number of flies within four body lengths (4BL, which corresponds to ∼1cm) of each individual. Each chamber constitutes one replicate, with a maximum of three replicates performed per genotype and sex per day, for a total of 7 to 15 replicates analyzed across multiple experimental days.

### Longevity assays

Longevity was assessed in both male and female flies for all experimental and control genotypes, as in Brenman et al. 2018 [52]. Same-sex flies of 3-4 days were housed in groups of 20 per vial containing standard food media and maintained at 25°C, approximately 50% relative humidity, under a 12:12 light-dark cycle. Flies were transferred to fresh food vials twice weekly, and the number of dead flies was recorded at each transfer. For each genotype and sex, 3 to 4 vials per cohort, and t3 to 5 cohorts were analyzed.

### Statistical analysis

Statistical analysis and graphical representation were performed in GraphPad Prism (RRID:SCR_002798, version 11 for Mac, GraphPad Software, Boston, MA, USA, www.graphpad.com). We first assessed the normality and homoscedasticity of the data distribution prior to applying the appropriate parametric tests (simple t-test or ANOVA or Welch t-test/ANOVA when the standard deviations were different among groups). When there were more than two groups but only one variable, we used a one-way ANOVA. One-way ANOVAs were followed by a post hoc test to identify significantly different points, while correcting for multiple comparisons. We used the Kaplan-Meier method to estimate survival probability, and Mantel-Cox log-rank to test differences between survival curves

## Results

We independently reanalyzed, using a slightly different methodology (see methods) the RNA-seq dataset published by Xie et al. (2023) to characterize transcriptional changes following loss of nlg3. Differentially expressed genes were identified using |log₂FC| ≥ 1.3 and p < 0.05. We identified 555 significantly differentially expressed genes in *nlg3* knockout flies compared with wild-type controls, including 342 upregulated and 213 downregulated genes (Fig. 1A). Among the most strongly upregulated genes were *Amy-d, Amy-p, AttC, CG15337, CG30098, CR41504, Diedel, DptA, Dro,* and *Ebp*. Several of these genes (*AttC, CG30098 Diedel, DptA,* and *Dro*) are associated with immune or stress-related responses [53, 54], consistent with the prominent transcriptional changes reported in the original study [35]. In our analysis, the most strongly downregulated genes included *Hsp70Ba, bard, CG13043, CG15480, CG18577, CG42681, CG43915, Elba3, Ir92a,* and *spo*. Of those, only the function of *Hsp70Ba, Elba3, Ir92a,* and *spo* are known, although *bard* has been associated to ubiquitin-dependent protein catabolic processes [55]. *Elba3 is involved in , Ir92a is, and spo.Hsp70Ba* encodes a member of the heat-shock protein 70 family, which contributes to thermotolerance and recovery from heat shock; Hsp70Ba expression is also induced by hypoxia in Drosophila cells [56, 57]. *Elba3* is involved in chromatin insulation and the regulation of gene expression as a component of the ELBA complex [58]; *Ir92a* encodes a chemosensory ionotropic receptor involved in amine detection and attraction behaviour [59]; and *spo (spook)* encodes the cytochrome P450 enzyme CYP307A1, which participates in ecdysone biosynthesis [60].

**Figure 1.**
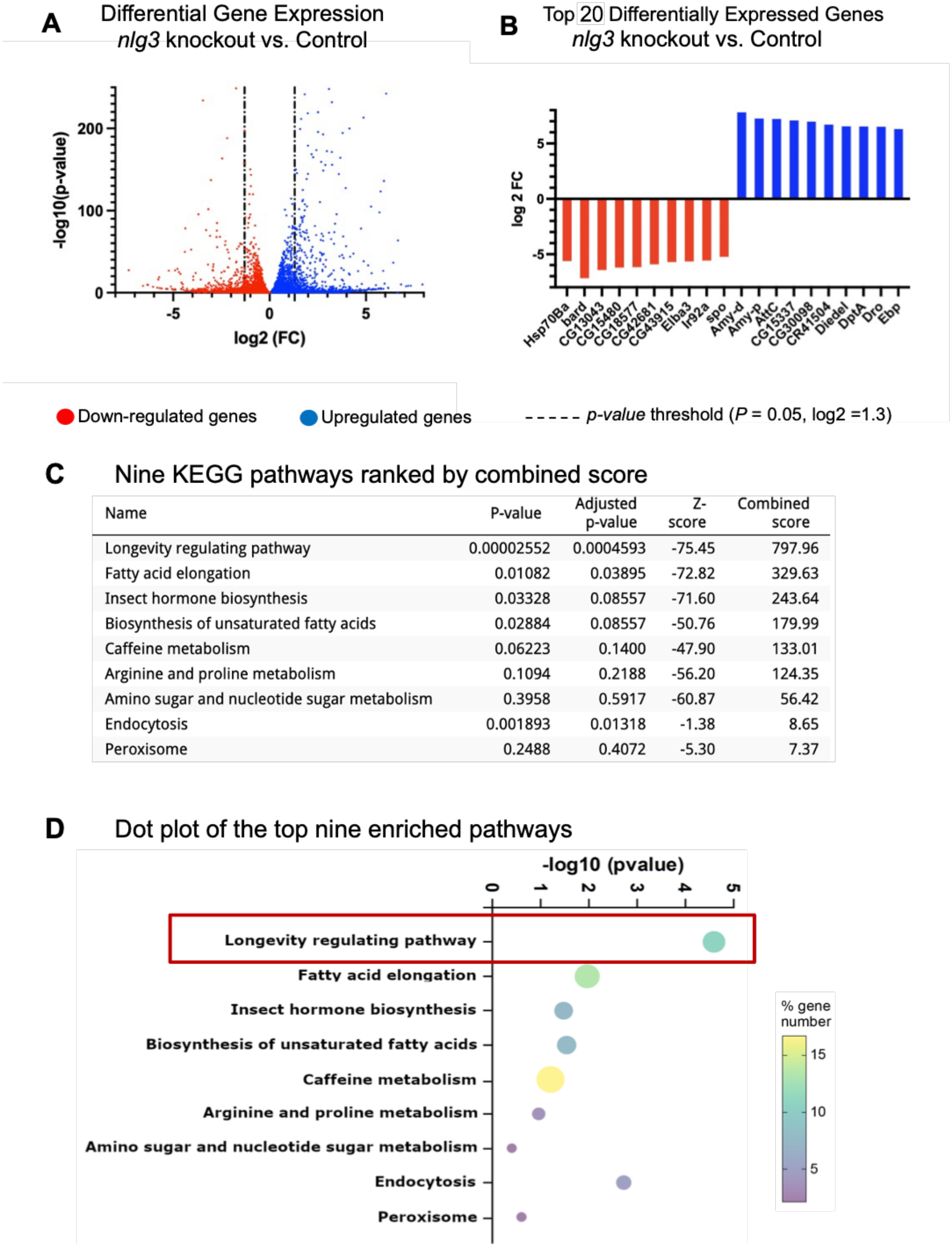
Transcriptome changes following *nlg3* loss of function. **(A)** Each dot represents a gene from the RNA-seq analysis comparing *nlg3* knockout flies to control flies. The x-axis shows the log₂ (fold change) in gene expression, and the y-axis shows the −log₁₀(*p*-value). Genes in blue are upregulated and genes in red are downregulated. Dashed lines indicate the differential expression thresholds: log₂FC ≤ −1.3 or ≥ 1.3, and *p* < 0.05. **(B)** Bar plot showing the top 20 most significantly differentially expressed genes in *nlg3* knockout flies compared to control flies, ranked by log₂(fold change). Red bars indicate downregulated genes, and blue bars indicate upregulated genes. **(C)** Table showing the top nine KEGG enriched pathways from the RNA-seq analysis comparing *nlg3* knockout flies to control flies. For each pathway, the p-value, adjusted *p*-value, *Z*-score, and combined score obtained from FlyEnrichr are shown. The longevity regulating pathway was the most significantly enriched pathway. **(D)** Dot plot of the top nine enriched pathways from the RNA-seq analysis comparing *nlg3* knockout flies to control flies. The x-axis shows the −log₁₀(*p*-value). The size and colour of each dot represent the percentage of genes in each pathway, as indicated by the colour scale. The longevity regulating pathway showed the strongest enrichment.

### KEGG pathway enrichment analysis

The overall distribution of differentially expressed genes was visualized using a volcano plot on DESeq2, with significantly upregulated and downregulated genes distinguished according to their direction of expression change (Fig. 1A). The most strongly altered genes were additionally highlighted to illustrate the major transcriptional changes associated with loss of *nlg3*. The ten most upregulated and ten most downregulated genes were further visualized according to their expression changes, also using DESeq2 (Fig. 1B). Together, our re-analysis confirmed the broad transcriptional response associated with *nlg3* loss while also identifying a prominent reduction in *Hsp70Ba* expression, a gene encoding a member of the heat-shock protein 70 family, which prompted further investigation of pathways associated with the downregulated gene set.

To investigate the biological pathways associated with the transcriptional changes, KEGG pathway enrichment analysis was performed separately on the downregulated genes using flyEnrichr. The longevity-regulating pathway was the most significantly enriched KEGG pathway in our analysis (p = 2.55 × 10⁻⁵; Fig. 1C-D). In addition to the longevity-regulating pathway, endocytosis and fatty acid elongation were significantly enriched after correction for multiple testing (adjusted *p* = 0.01318 and 0.03895, respectively). Insect hormone biosynthesis and biosynthesis of unsaturated fatty acids showed nominal enrichment (*p* < 0.05), but neither remained significant after correction (adjusted *p* = 0.08557 for both).

The enrichment of the longevity-regulating pathway was associated with the downregulation of six heat-shock protein genes: *Hsp68, Hsp70Ba, Hsp70Ab, Hsp70Bc, Hsp70Bb,* and *Hsp70Aa*. Notably, the same group of heat-shock protein genes also contributed to the enrichment of the endocytosis, spliceosome, and protein-processing pathways. In addition, *Elo68alpha* and *Elo68beta* were associated with fatty acid elongation and unsaturated fatty acid biosynthesis, while *spo* and *spok* contributed to the enrichment of the insect hormone biosynthesis pathway. Together, these results identify the heat-shock protein genes as major contributors to the pathway enrichment observed among the downregulated genes.

### RT-qPCR validation of reduced *hsp68* and *hsp70* expression in *nlg3* mutants

To validate the transcriptional changes identified by RNA-seq and subsequent pathway enrichment analysis, RT-qPCR was performed to examine the expression of heat-shock protein genes in *nlg3* mutant flies. *hsp68* and *hsp70* expression was assessed in Cs (+/+), *nlg3* heterozygous (*nlg3^def^/*+), and *nlg3* deficiency (*nlg3^def^/ nlg3^def^*) fly heads. Consistent with the RNA-seq results, expression of *hsp70* was reduced in the heterozygous genotype and both *hsp68* and *hsp70* showed a reduction in the *nlg3* deficiency compared with controls (Fig. 2). Due to the high sequence similarity among the *hsp70* paralogs identified in the RNA-seq analysis (*hsp70Aa, hsp70Ab, hsp70Ba, hsp70Bb,* and *hsp70Bc*), paralog-specific primer pairs could not be reliably designed. Therefore, RT-qPCR was used to assess *hsp70* expression collectively rather than to quantify individual *hsp70* paralogs.

**Figure 2.**
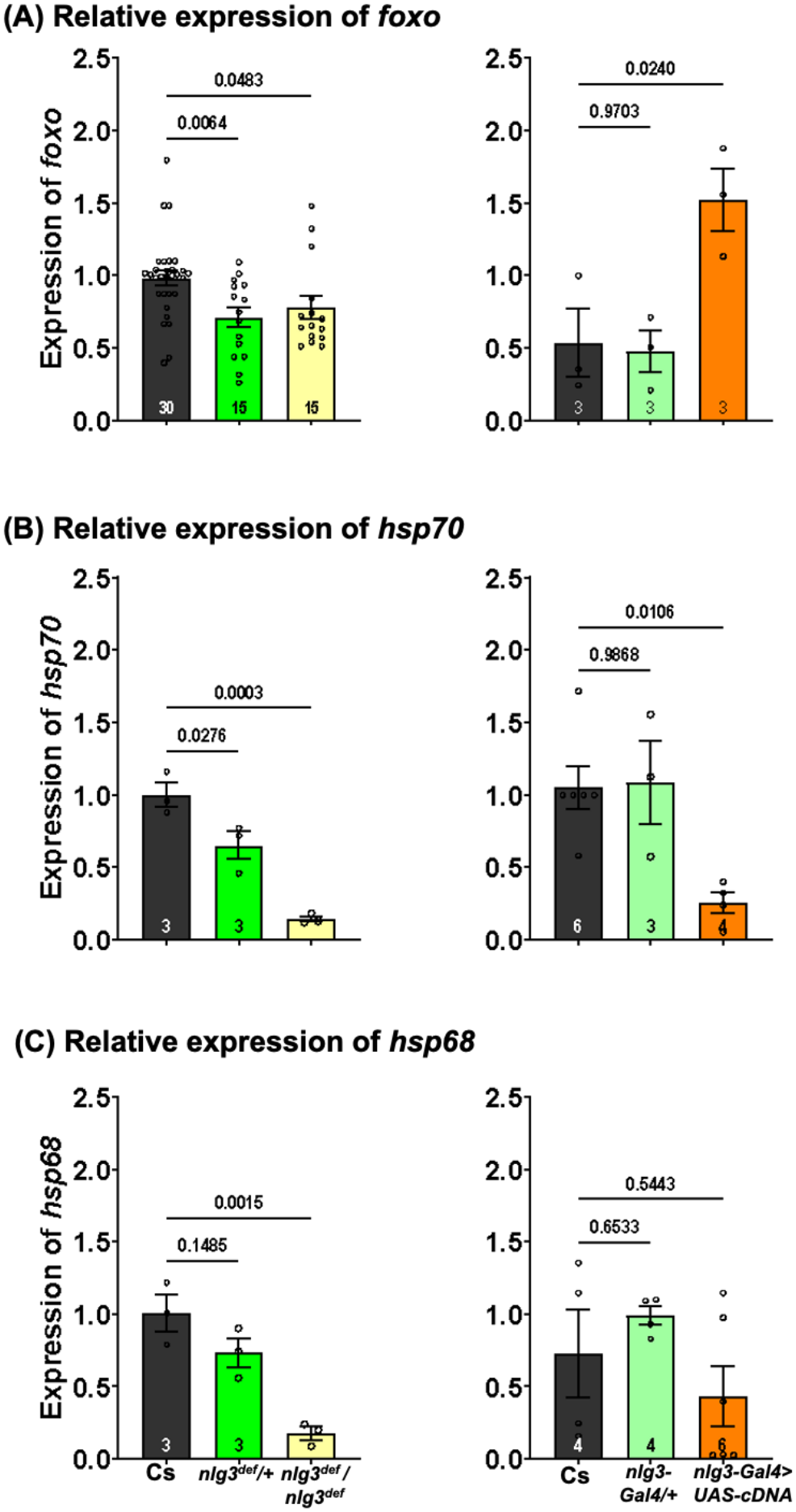
*Neuroligin 3* deficiency and overexpression differentially alter longevity-associated transcripts. Relative expressions of **(A)** *foxo*, **(B)** *hsp70*, and **(C)** *hsp68* were quantified by RT-qPCR in adult fly heads. Left panels show heterozygous (*nlg3^def^*/+) and homozygous (*nlg3^def^*/*nlg3^def^*) deficiency genotypes, while right panels show *nlg3* overexpression (*nlg3*-*Gal4>UAS-cDNA^nlg3^*) and *nlg3-Gal4* driver only (*nlg3*-*Gal4*/+). All comparisons were performed against Canton-S (Cs). Relative transcript levels were calculated using the ΔΔCt method, normalized to *αTub84B*. Bars represent means ± SEM, individual dots indicate plotted observations, and numbers within bars indicate sample sizes. Data were analyzed using separate ordinary one-way ANOVAs for deficiency and overexpression datasets, followed by Dunnett’s multiple-comparisons tests. **(A)** *foxo* expression differed among deficiency genotypes (*F*(2, 57) = 5.628, p = 0.0059), with reduced expression in both heterozygous (Dunnett’s post hoc p = 0.0064) and homozygous (Dunnett’s post hoc p = 0.0483) deficiency flies. Expression also differed among overexpression genotypes (*F*(2, 6) = 8.435, p = 0.0181), with increased expression following *nlg3* overexpression (Dunnett’s post hoc p = 0.0240), but no detectable difference in *nlg3-Gal4* driver/+ (Dunnett’s post hoc p = 0.9703). **(B)** *hsp70* expression differed among deficiency genotypes (*F*(2, 6) = 33.71, p = 0.0005), with reduced expression in heterozygous (Dunnett’s post hoc p = 0.0276) and homozygous (Dunnett’s post hoc p = 0.0003) deficiency flies. Expression also differed among overexpression genotypes (*F*(2, 6) = 7.334, p = 0.0109), with reduced expression following *nlg3* overexpression (Dunnett’s post hoc p = 0.0106), but no detectable difference in *nlg3*-Gal4 driver/+ (Dunnett’s post hoc p = 0.9868). **(C)** *hsp68* expression differed among deficiency genotypes (*F*(2, 6) = 19.83, p = 0.0023), with reduced expression in homozygous (Dunnett’s post hoc p = 0.0015), but not heterozygous (Dunnett’s post hoc p = 0.1485), deficiency flies. No differences were detected among overexpression genotypes (*F*(2, 11) = 1.675, p = 0.2317), with neither *nlg3* overexpression (Dunnett’s post hoc, p = 0.5443) nor *nlg3-Gal4* driver/+ (Dunnett’s post hoc, p = 0.6533) differing from Cs.

### Loss of *neuroligin 3* reduces stress-response transcripts, whereas overexpression differentially affects *hsp70* and *foxo*

FOXO (Forkhead box O) is an important contributor to longevity across species[61], that appears on the KEGG longevity-regulating pathways[62]. This transcription factor also regulates the expression of stress-response genes, including heat-shock proteins [63]. We thus next examined whether *foxo* expression was altered following loss of *nlg3*. Expression of *foxo* was reduced in both heterozygous and homozygous deficiency flies compared with Cs (Ordinary One-Way ANOVA; *F*(2, 57) = 5.628, Dunnett’s post-hoc p = 0.0059; *nlg3*^def^/+, p = 0.0064; *nlg3*^def^/*nlg3*^def^, p *=* 0.0483; Fig. 2A). In contrast, *nlg3* overexpression was associated with increased *foxo* expression relative to control (Ordinary One-Way ANOVA; *F* (2, 6) = 8.435, Dunnett’s post-hoc p = 0.0181; *nlg3-Trojan-Gal4>UAS-cDNA^nlg3^*, p = 0.024), whereas the heterozygote *nlg3-*Trojan-Gal4 driver did not differ from Cs (*nlg3*-*Trojan-*Gal4/+, Dunnett’s post-hoc p = 0.9703). Expression of *hsp70* was reduced in both heterozygous and homozygous deficiency flies compared with control (Ordinary One-Way ANOVA; *F* (2, 6) = 33.71, p = 0.0005; Dunnett’s post-hoc *nlg3*^def^/*+*, p = 0.0276; *nlg3*^def^/*nlg3^def^,* p = 0.0003), with a greater reduction observed in homozygous deficiency flies (Fig. 2B). Similarly, *nlg3* overexpression was associated with lower *hsp70* expression compared with Cs (Ordinary One-Way ANOVA; *F* (2, 6) = 7.334, p = 0.0109; Dunnett’s post-hoc *nlg3-Trojan-Gal4>UAS-cDNA^nlg3^*, p = 0.0106), while the heterozygote *nlg3-*Trojan-Gal4 driver showed no detectable difference (p = 0.9868).

Expression of *hsp68* was reduced in homozygous deficiency flies compared with Cs (Ordinary One-Way ANOVA; *F* (2, 6) = 19.83, p = 0.0023; Dunnett’s post-hoc *nlg3*^def^/*nlg3^def^, p* = 0.0015), whereas heterozygous deficiency did not produce any difference (Dunnett’s post-hoc p = 0.1485; Fig. 2C). Neither *nlg3* overexpression nor the *nlg3-*Trojan-Gal4 driver/+ differed from Cs (Ordinary One-Way ANOVA; *F* (2, 11) = 1.675, p = 0.2317; Dunnett’s post-hoc *nlg3-Trojan-Gal4>UAS-cDNA^nlg3^*, p = 0.5443; nlg3*-Trojan-Gal4*/+, p = 0.6533).

### *Neuroligin 3* overexpression shortens lifespan, while heterozygous deficiency extends lifespan in both sexes

Because of the enrichment of the longevity-regulating pathway and the changes in stress-response gene expression, we next examined whether manipulation of *nlg3* was associated with altered organismal longevity. To investigate whether *nlg3* influences organismal longevity, survival was examined in male and female *Drosophila* carrying heterozygous or homozygous *nlg3* deficiencies, *RNAi*-mediated knockdown, or *nlg3* overexpression, alongside their respective controls and *nlg3-*Trojan-Gal4 driver/+ (Fig. 3). The experiment was independently repeated 17 times; representative survival curves are shown. Kaplan–Meier survival analysis for the representative data revealed differences among genotypes in both males (Mantel–Cox test, χ²(7) = 263, *p* < 0.0001) and females (χ²(7) = 291.9, *p* < 0.0001; Fig. 3A). To further characterize these survival differences, total median lifespan and maximum longevity of all 17 replicates were compared among genotypes. Welch’s one-way ANOVA revealed differences in median lifespan in both males (*W*(7, 44.30) = 61.16, *p* < 0.0001; Fig. 3B) and females (*W*(7, 43.75) = 39.60, *p* < 0.0001; Fig. 3B). Similarly, maximum longevity differed among genotypes in males (*W*(7, 103.5) = 77.41, *p* < 0.0001; Fig. 3C) and females (*W*(7, 45.12) = 171.12, *p* < 0.0001; Fig. 3C). Median survival differed among *nlg3* genotypes in male *Drosophila* (Fig. 3B). Relative to Cs control, *nlg3* overexpression reduced median survival by 27% (Dunnett’s T3, *p* < 0.0001). Homozygous *nlg3* deficiency produced a reduction of 28% (*p* < 0.0001). No difference was detected between the *nlg3* overexpression and *nlg3* deletion genotypes (*p* > 0.9999). In contrast, heterozygous *nlg3* deficiency increased median survival by 29.5% relative to Cs (*p* = 0.0001). The *nlg3-Gal4* driver/+ and *RNAi*-mediated knockdown also exhibited increased median survival of 21.6% (*p* = 0.0002) and 31.6% (*p* < 0.0001), respectively. However, median survival did not differ between *RNAi*-mediated knockdown and the *nlg3-Gal4* driver/+ (*p* = 0.4164), or between *RNAi*-mediated knockdown and heterozygous deficiency (*p* > 0.9999).

**Fig. 3.**
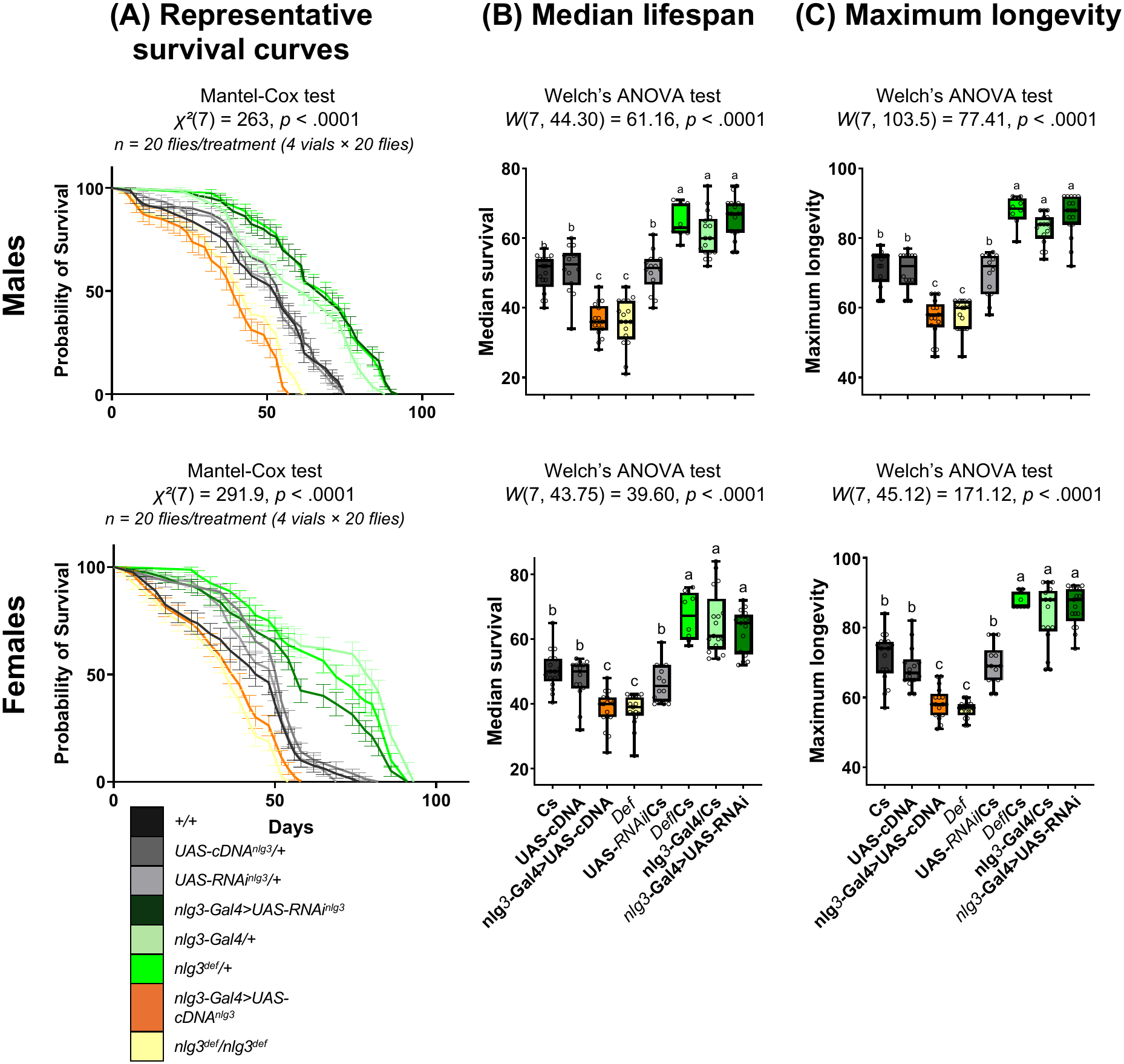
Survival and longevity following manipulation of *nlg3* in male and female *Drosophila melanogaster*. Survival curves **(A)**, median lifespan **(B)**, and maximum longevity **(C)** are shown for males (top row) and females (bottom row). Eight genotypes were examined, including Canton-S (Cs), *nlg3*-Gal4/+, *nlg3-Gal4>UAS-RNAi^nlg3^*, *UAS-RNAi^nlg3^/*+, *nlg3-Gal4>UAS-cDNA^nlg3^*, *UAS-cDNA^nlg3^/+*, homozygous *nlg3* deficiency (*nlg3^def^*), and heterozygous deficiency (*nlg3^def^/+*). Each genotype consisted of 17 replicate vials containing 20 same-sex flies (*n* = 340 flies per genotype per sex). Representative survival curves are shown, containing 4 replicate vials of 20 flies per treatment (n = 20 flies per genotype per sex). Gray represents control genotypes, orange represents *nlg3* overexpression, yellow represents homozygous deficiency, and green represents genotypes with reduced *nlg3* function. **(A)** Survival curves illustrate differences in survival probability across genotypes. Survival distributions differed among genotypes in males (log-rank Mantel–Cox test, χ²(7) = 1063, *p* < 0.0001) and females (χ²(7) = 900.4, *p* < 0.0001). Error bars represent standard error. **(B)** Median lifespan was determined for each replicate vial and compared among genotypes using Welch’s one-way ANOVA. Differences were detected in both males (*W*(7, 44.30) = 61.16, *p* < 0.0001) and females (*W*(7, 43.75) = 39.60, *p* < 0.0001). **(C)** Maximum longevity, defined as the greatest recorded lifespan within each replicate vial, also differed among genotypes in males (*W*(7, 103.5) = 77.41, *p* < 0.0001) and females (*W*(7, 45.12) = 171.12, *p* < 0.0001). For panels B and C, each point represents one replicate vial. Box plots display the median and interquartile range (IQR), with whiskers extending to the minimum and maximum observed values. All individual data points are shown. Welch’s ANOVAs were followed by Dunnett’s T3 multiple-comparisons tests. Different letters indicate statistical differences among genotypes (*p* < 0.05).

**Figure 4.**
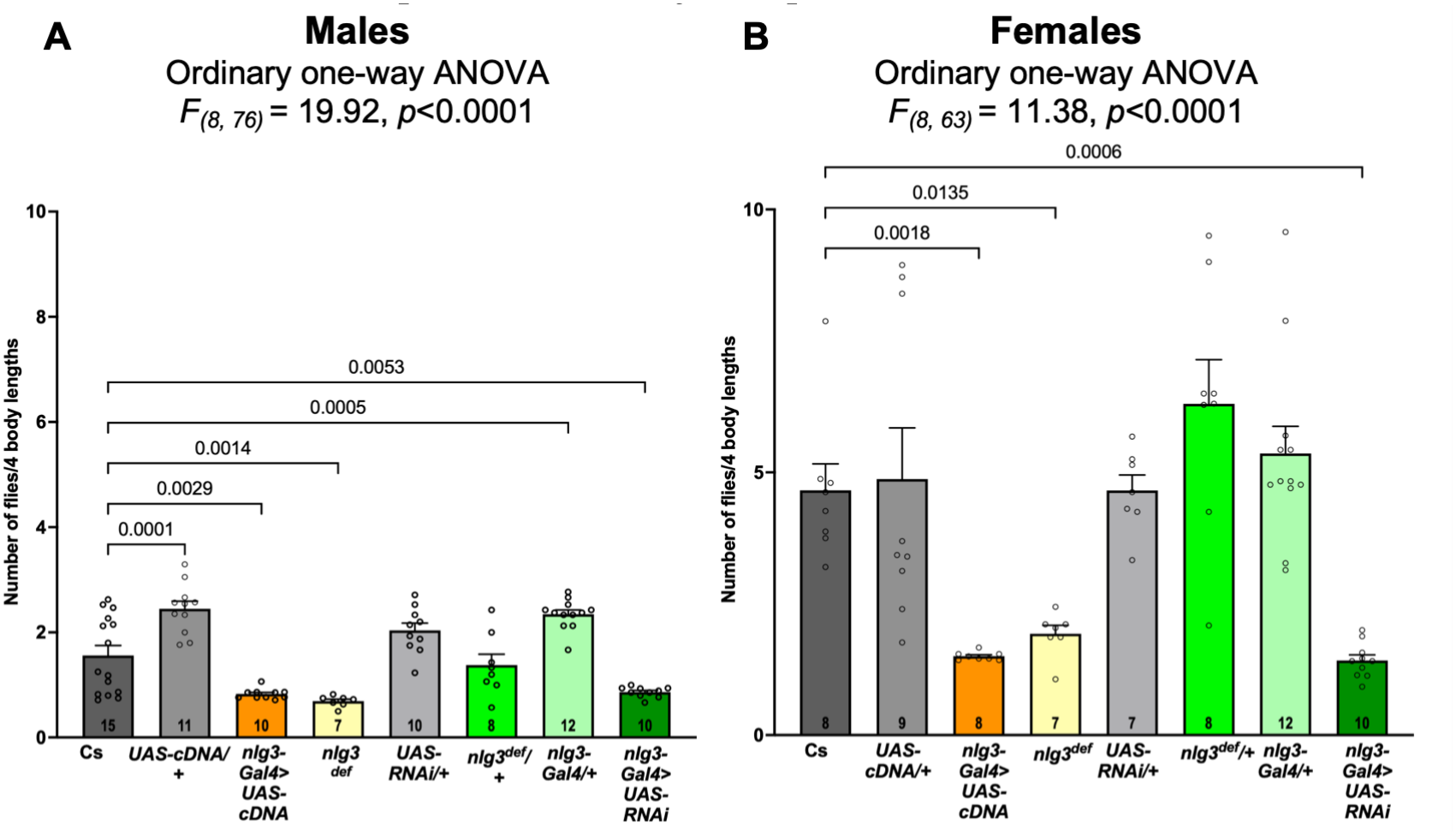
Manipulation of *nlg3* alters social spacing in male and female *Drosophila melanogaster*. Bars show the mean number of flies within four body lengths (± SEM), with lower values indicating greater social spacing. Each dot represents one observation, and numbers within bars indicate sample sizes. Genotypes are indicated below each bar. Data were analyzed separately for each sex using one-way ANOVA (males: *F*(8, 76) = 19.92, *p* < 0.0001; females: *F*(8, 63) = 11.38, *p* < 0.0001), followed by Šídák’s multiple-comparisons test to control. Brackets show selected comparisons with adjusted *p* values.

In females, median survival differed among *nlg3* genotypes (Fig. 3B). Relative to Cs control, *nlg3* overexpression reduced median survival by 23.8% (Dunnett’s T3, *p* < 0.0001). Homozygous *nlg3* deficiency produced a similar reduction of 24.5% (*p* < 0.0001). No difference was detected between the *nlg3* overexpression and *nlg3* deletion genotypes (*p* > 0.9999). In contrast, heterozygous *nlg3* deficiency increased median survival by 32.8% relative to Cs (*p* = 0.0001). The *nlg3-Gal4*/+ driver control and *RNAi*-mediated knockdown also exhibited increased median survival of 28.9% (*p* = 0.0002) and 23% (*p* < 0.0001), respectively. However, median survival did not differ between *RNAi*-mediated knockdown and the *nlg3-Gal4*/+ driver control (*p* = 0.4164), or between *RNAi*-mediated knockdown and heterozygous deficiency (*p* > 0.9999). Genetic manipulation of *nlg3* produced distinct effects on maximum recorded lifespan in male and female *Drosophila* (Fig. 3C). Relative to Cs, *nlg3* overexpression was associated with reductions of 17.95% in males and 21.43% in females, while homozygous *nlg3* deficiency showed reductions of 20.51% and 28.57%, respectively. However, the *nlg3*-Gal4/+ hypomorphic genotype exhibited increases of 12.82% in males and 10.71% in females. *RNAi*-mediated knockdown was similarly associated with increases of 17.95% and 9.52%, respectively. Heterozygous *nlg3* deficiency also showed increased maximum recorded lifespan, with increases of 17.95% in males and 8.33% in females relative to Cs control (see also **Table 3**).

**Table 2.**
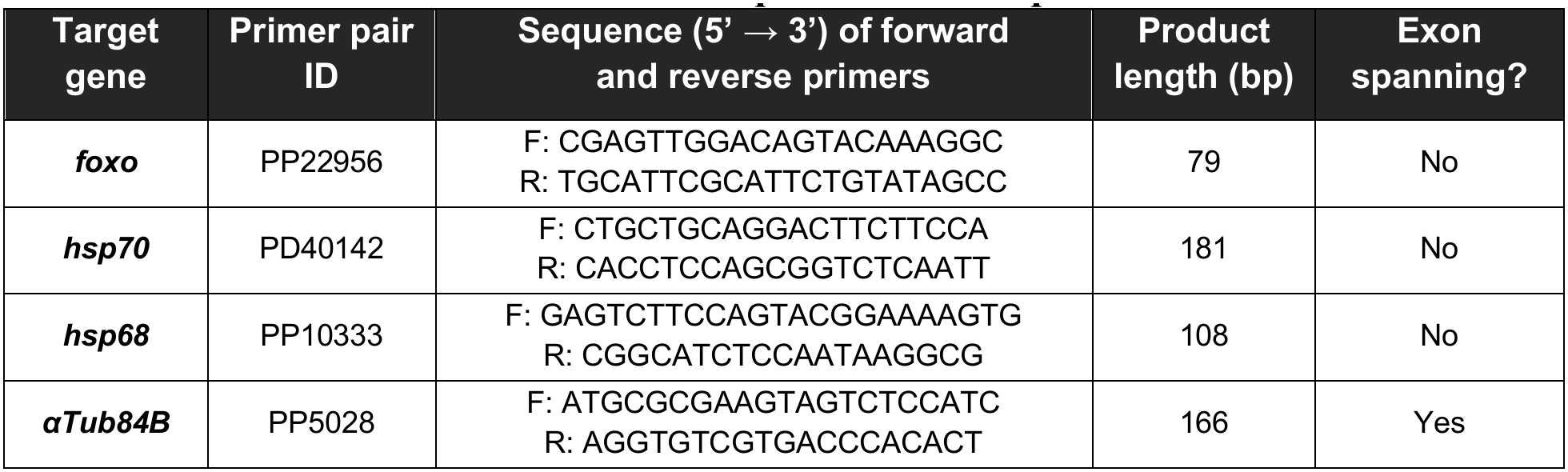
Primer sequences for RT-qPCR.

**Table 3.** Pairwise comparisons of median survival and maximum longevity following genetic manipulation of *nlg3* in male and female *Drosophila*. Median survival and maximum longevity were compared among Canton-S (Cs), *nlg3*-*Gal4*/+, *nlg3*-*Gal4>UAS-RNAi^nlg3^*, *nlg3-Gal4>UAS-cDNA^nlg3^*, homozygous *nlg3* deficiency (*nlg3^def^*), and heterozygous deficiency (*nlg3^def^/+*) genotypes. Analyses were performed separately for males and females using Welch’s one-way ANOVA, followed by Dunnett’s T3 multiple-comparisons tests to account for unequal variances. Mean differences are expressed in days and calculated as the mean of the first genotype minus that of the second genotype listed in each comparison. Positive values indicate greater median survival or maximum longevity in the first genotype, whereas negative values indicate lower values. The 95% confidence intervals (CI) and adjusted *p*-values are reported for each comparison.

|  | Dunnett's T3 multiple comparisons tests | Median survival |  |  | Maximum longevity |  |  |
|---|---|---|---|---|---|---|---|
|  |  | Mean diff. | 95.00% CI of diff. | Adjusted P Value | Mean diff. | 95.00% CI of diff. | Adjusted P Value |
| Males | *Cs vs. nlg3-Gal4/+* | -10.82 | -17.64 to -4.011 | 0.0002 | -11.12 | -16.77 to -5.462 | <0.0001 |
|  | *Cs vs. nlg3-Gal4>UAS-RNAi* | -15.79 | -22.10 to -9.488 | <0.0001 | -14.94 | -21.47 to -8.409 | <0.0001 |
|  | *Cs vs. nlg3-Gal4>UAS-cDNA* | 13.59 | 7.817 to 19.36 | <0.0001 | 14.65 | 8.337 to 20.96 | <0.0001 |
|  | *Cs vs. Def* | 14.18 | 7.029 to 21.32 | <0.0001 | 13.82 | 8.052 to 19.60 | <0.0001 |
|  | *Cs vs. Def/+* | -14.75 | -22.69 to -6.809 | 0.0001 | -16.1 | -23.30 to -8.901 | <0.0001 |
|  | *nlg3-Gal4/+ vs. nlg3-Gal4>UAS-RNAi* | -4.971 | -11.97 to 2.032 | 0.4164 | -3.824 | -9.860 to 2.213 | 0.6003 |
|  | *nlg3-Gal4/+ vs. nlg3-Gal4>UAS-cDNA* | 24.41 | 17.85 to 30.97 | <0.0001 | 25.76 | 19.99 to 31.54 | <0.0001 |
|  | *nlg3-Gal4/+ vs. Def* | 25 | 17.28 to 32.72 | <0.0001 | 24.94 | 19.81 to 30.07 | <0.0001 |
|  | *nlg3-Gal4/+ vs. Def/+* | -3.926 | -12.29 to 4.434 | 0.8935 | -4.985 | -11.83 to 1.860 | 0.288 |
|  | *nlg3-Gal4>UAS-RNAi vs. nlg3-Gal4>UAS-cDNA* | 29.38 | 23.37 to 35.39 | <0.0001 | 29.59 | 22.97 to 36.21 | <0.0001 |
|  | *nlg3-Gal4>UAS-RNAi vs. Def* | 29.97 | 22.66 to 37.28 | <0.0001 | 28.76 | 22.64 to 34.89 | <0.0001 |
|  | *nlg3-Gal4>UAS-RNAi vs. Def/+* | 1.044 | -7.008 to 9.096 | >0.9999 | -1.162 | -8.563 to 6.240 | >0.9999 |
|  | *nlg3-Gal4>UAS-cDNA vs. Def* | 0.5882 |  | >0.9999 | -0.8235 | -6.697 to 5.050 | >0.9999 |
|  | *nlg3-Gal4>UAS-cDNA vs. Def/+* | -28.34 | -36.18 to -20.50 | <0.0001 | -30.75 | -37.99 to -23.51 | <0.0001 |
|  | *Def vs. Def/+* | -28.93 | -37.51 to -20.34 | <0.0001 | -29.93 | -36.81 to -23.05 | <0.0001 |
| Females | *Cs vs. nlg3-Gal4/+* | -10.82 | -17.64 to -4.011 | 0.0002 | 15.29 | 8.882 to 21.71 | <0.0001 |
|  | *Cs vs. nlg3-Gal4>UAS-RNAi* | -15.79 | -22.10 to -9.488 | <0.0001 | 13.29 | 6.377 to 20.21 | <0.0001 |
|  | *Cs vs. nlg3-Gal4>UAS-cDNA* | 13.59 | 7.817 to 19.36 | <0.0001 | -14.41 | -21.82 to -7.000 | <0.0001 |
|  | *Cs vs. Def* | 14.18 | 7.029 to 21.32 | <0.0001 | -15.74 | -22.40 to -9.073 | <0.0001 |
|  | *Cs vs. Def/+* | -14.75 | -22.69 to -6.809 | 0.0001 | -11.88 | -21.03 to -2.738 | 0.0033 |
|  | *nlg3-Gal4/+ vs. nlg3-Gal4>UAS-RNAi* | -4.971 | -11.97 to 2.032 | 0.4164 | -2 | -6.183 to 2.183 | 0.9169 |
|  | *nlg3-Gal4/+ vs. nlg3-Gal4>UAS-cDNA* | 24.41 | 17.85 to 30.97 | <0.0001 | -29.71 | -34.85 to -24.56 | <0.0001 |
|  | *nlg3-Gal4/+ vs. Def* | 25 | 17.28 to 32.72 | <0.0001 | -31.03 | -34.68 to -27.38 | <0.0001 |
|  | *nlg3-Gal4/+ vs. Def/+* | -3.926 | -12.29 to 4.434 | 0.8935 | -27.18 | -34.94 to -19.41 | <0.0001 |
|  | *nlg3-Gal4>UAS-RNAi vs. UAS-RNAi/+* | 15.22 | 8.196 to 22.25 | <0.0001 | -27.71 | -33.54 to -21.87 | <0.0001 |
|  | *nlg3-Gal4>UAS-RNAi vs. nlg3-Gal4>UAS-cDNA* | 29.38 | 23.37 to 35.39 | <0.0001 | -29.03 | -33.66 to -24.39 | <0.0001 |
|  | *nlg3-Gal4>UAS-RNAi vs. Def/+* | 1.044 | -7.008 to 9.096 | >0.9999 | -25.18 | -33.29 to -17.06 | <0.0001 |
|  | *nlg3-Gal4>UAS-cDNA vs. Def* | 0.5882 |  | >0.9999 | -1.324 | -6.787 to 4.140 | >0.9999 |
|  | *nlg3-Gal4>UAS-cDNA vs. Def/+* | -28.34 | -36.18 to -20.50 | <0.0001 | 2.529 | -5.982 to 11.04 | 0.9997 |
|  | *Def vs. Def/+* | -28.93 | -37.51 to -20.34 | <0.0001 | 3.853 | -4.052 to 11.76 | 0.8783 |

### Overexpressing *neuroligin 3* increases social space between neighbouring flies

We reported that loss of function in *nlg3^def^* and *RNAi*-mediated knockdown using *nlg3*-Trojan-*Gal4* lead to reduction of number of flies within 4 body length (4BL), showing an increased social spacing [32]. Indeed, Lower values of 4BL indicate that flies settle farther apart, corresponding to increased social spacing. To examine whether graded changes in *nlg3* influence social behaviour, we measured the number of flies within 4BL in the same genotypes as those tested for longevity. In males, social spacing differed among genotypes (Ordinary one-way ANOVA, *F*_(8, 76)_ = 19.92, *p* < 0.0001; Fig. 5): as previously reported by Robinson et al., 2023 [32], relative to Cs, homozygous deficiency (*nlg3^def^*) increased spacing (Šídák post-hoc p = 0.0014), when heterozygous for the deficiency did not show altered social space. Also, as previously reported, *RNAi*-mediated knockdown with heterozygous *nlg3*-Trojan-*Gal4* led to flies further apart (*nlg3-Gal4/UAS-RNA^nlg3^*; Šídák post-hoc p = 0.0053). Males were also further apart when *nlg3* was overexpressed (*nlg3-Gal4/UAS-cDNA^nlg3^*; Šídák post-hoc p = 0.0029). Of note, the hypomorph heterozygous *nlg3*-Trojan-*Gal4*/+ males were closer compared to Cs, as were the *UAS-cDNA^nlg3^/+* lines (*nlg3-Gal4/+,* Šídák p = 0.0005; *UAS-cDNA^nlg3^*/+, Šídák p = 0.0001). In females, social spacing also differed among genotypes (Ordinary one-way ANOVA, *F*_(8, 63)_ = 11.38, *p* < 0.0001). And as previously reported, both the deficiency and *RNAi*-mediated knockdown of *nlg3* led to flies being further apart relative to Cs (*nlg3^def^*, Šídák p = 0.0135; *nlg3-Gal4/UAS-RNAi*; Šídák p = 0.0006). Similarly to those genotypes and male flies, overexpression of *nlg3* decreased the number of flies within four body lengths, indicating increased social spacing (Šídák; p = 0.0018). None of the other genotypes altered social spacing in females.

## Discussion

Our findings identify a previously unrecognized role for *nlg3* in the regulation of lifespan in *Drosophila melanogaster*. Reanalysis of the publicly available *nlg3* knockout RNA-seq dataset provided a way to identify additional changes relevant to the longevity phenotypes examined here. Although our analysis identified a different number of differentially expressed genes than the original study, it recovered a prominent upregulation of immune-associated genes, *AttC, CG30098 Diedel, DptA,* and *Dro*. This general agreement supports the conclusion that loss of *nlg3* alters innate immune-related transcription in adult fly heads. Differences in the number and identity of significant genes may reflect differences in the analytical workflow and statistical criteria used for reanalysis; therefore, individual gene-level differences should be interpreted cautiously. The immune-associated signature provides context for the broad transcriptional effects of *nlg3* loss, while the downregulated genes offered a complementary avenue for investigating pathways potentially related to longevity. Of note, one of the GO enrichment pathway reported by Xie et al. (2023) [35] is related to longevity: the heat-shock-mediated protein folding and stress response pathway to heat and hypoxia also were identified among the down-regulated genes. Hence, both approaches validate each other.

Our analysis of downregulated genes identified enrichment of the KEGG longevity-regulating pathway, driven by several *Hsp70* family genes and *Hsp68*. RT-qPCR in *nlg3* mutant flies supported reduced expression of *Hsp68* and *Hsp70s*, as well as reduced *foxo* expression. These findings are consistent with altered expression of FOXO-associated stress-response genes following *nlg3* loss and suggest a potential link between *neuroligin*-dependent transcriptional regulation and longevity-related pathways. However, reduced *foxo* transcript abundance does not establish a change in FOXO protein activity, and the collective *Hsp70* assay does not resolve expression of individual paralogs. Further experiments will be needed to determine whether changes in FOXO activity or heat-shock protein expression contribute causally to the observed longevity phenotypes.

Partial and complete *nlg3* deficiency produced distinct behavioural and longevity outcomes. It is important to note that we used as a driver of expression in *nlg3*-expressing cell a Trojan insertion in the 3^rd^ intron of the *nlg3* gene. Trojan exons are designed to truncate the gene host transcript [64], so *nlg3-Trojan-Gal4/+* is a heterozygous loss-of-function. We have indeed shown that there are reduced levels of Nlg3 protein in both heterozygote and homozygote flies carrying *nlg3-Trojan-Gal4* (data not shown). Furthermore, we cannot obtain enough homozygote flies carrying *nlg3-Trojan-Gal4* to perform behaviour or longevity experiments, as those flies appear sublethal at the adult stage. This hints at possibility complex regulatory mechanisms for the expression and role of the *nlg3* gene and Nlg3 protein: indeed, the complete deficiency of *nlg3* is viable, although with a longevity reduction, but with increased fertility (data not shown).So, a homozygous insertion of a Trojan construct leads to some protein still being produced (data not shown), but strongly affected survival to adulthood, while removing the whole gene sequence does not prevent high fertility (data not shown). Some of the differences between the two types of manipulations are worth noting: besides *nlg3*, a few other small-size genes are also removed by the deficiency (*αEst-1*, *αEst-2*, *αEst-3* and *αEst-4*) [65], as well as the lncRNA included in the first large intron of *nlg3* (lncRNA:CR44332). The effects observed in the homozygous deficiency flies could be also the result of missing those genes. In contrast, the Trojan insertion does not remove lncRNA:CR44332, which might have some regulatory roles on *nlg3* expression itself. But this interrupted Nlg3 peptide might have some negative gain of function, leading to the reduced survival observed. Finally, rescue experiments will not be straightforward. Using a pan-neuronal driver, we have previously shown that the *UAS-cDNA^nlg3^* is expressed at high levels [51], which makes reaching the ideal dosage of *nlg3* possibly difficult. To address some of the limitations of the current genotypes tested, future work includes testing a trans-heterozygote (*nlg3^Def^ /nlg3-Trojan Gal4*) and a *cDNA^nlg3^*rescue in that *nlg3^Def^ /nlg3-Trojan Gal4* background. However, we can report the following: one-copy reduction of the *nlg3* gene (*nlg3^Def^/+, nlg3 Trojan Gal4/+*) leads to long-lived flies, with no additional effect of the RNAi*^nlg3^* compared its driver *nlg3 Trojan Gal4/+*. Complete loss (*Def/Def*) and overexpression of *nlg3* leads to equally short-lived flies. The UAS/+ lines have the same longevity as the Cs, to which they been outcrossed.

In contrast, one-copy reduction of the *nlg3* gene (*nlg3^Def^/+, nlg3 Trojan Gal4/+*) has either no effect on social spacing, or a reduced social spacing for males for *nlg3 Trojan Gal4/+*. But now, driving the RNAi*^nlg3^* using the driver *nlg3 Trojan Gal4/+* leads to a strong increase in social spacing, in both sexes, as does the overexpression. This is another non-monotonic dosage response but based on different dosages. Now, we also observe, in males only, an effect of the UAS/+ lines, suggesting a possible leaky effect, to be confirm by RT-qPCR.

What we can conclude is that longevity and social spacing are traits that differ in their sensitivity to reduced Nlg3 protein function; partial reduction may retain sufficient function for typical social spacing while modifying processes relevant to survival. More extensive loss may compromise functions required for both outcomes.

The shortening of lifespan following overexpression further suggests that the relationship between *nlg3* abundance and survival may be nonlinear. A comparable non-monotonic dosage response has been demonstrated for *Drosophila Sir2*, for which moderate overexpression extended lifespan whereas high expression was detrimental [66]. However, establishing such a relationship for *nlg3* will require direct measurements of expression across genotypes, which still needs to be performed.

The differing effects of *nlg3* manipulation have on longevity, and social spacing suggests that these traits are not coupled. Overexpression and homozygous deficiency both increased spacing, while heterozygous deficiency extended lifespan without a detectable spacing change. Similar outcomes may therefore arise through different cellular perturbations, and altered spacing alone does not explain the survival differences. These comparisons favour a model in which the consequences of *nlg3* manipulation depend on the trait, genetic context, and extent of functional disruption, rather than on a uniform effect of reduced expression. Our RNAseq and RT-qPCR data further show that nlg3 loss-of-function downregulates the longevity-associated genes *foxo*, *hsp70*s, and *hsp68*.

As far as we know, Drosophila *dnlg3* studies to date tested synaptic and circuit phenotypes only, not relation to FOXO, Hsp70/Hsp68, or aging. The proteostasis side of our observation has independent, if indirect, precedent. Drosophila dFOXO directly transactivates Hsp70-family and small heat-shock genes under oxidative stress [67]. Additionally, the neuronal PI3K-Akt/IGF-1R signalling (acting as a master regulator of neurodevelopment, structural plasticity, and metabolic maintenance) can suppress FOXO1 transcription via a FOXO-binding site in its own promoter [68]. Thus, FOXO drives the expression of Hsp70/Hsp68, and neuronal signalling can tune FOXO activity [68]. So far, the only known link between a *nlg* and a stress-related longevity pathway comes from *C. elegans*: oxidative stress activates the transcription factor SKN-1/Nrf2, which increases levels of the only Nlg protein in the worm, NLG-1, at synapses, and the amount of NLG-1 in turn affects survival under oxidative stress [69,70]. This SKN-1/Nrf2 pathway operates separately from the DAF-16/FOXO pathway; both are inhibited by insulin/IGF-1 signalling and share some effects, but can be genetically separated. For example, heat-shock proteins are a FOXO output, not an SKN-1/Nrf2 output [70–72]. Our work is the first, as far as we know, that is linking a synaptic cell-adhesion gene to the FOXO-HSP axis specifically, in any organism.

## Conclusion

*dnlg3* is already recognized as a dosage-sensitive regulator of social spacing, but its role in aging has remained unexplored. We now link behavioural output to the Drosophila FOXO-heat-shock-protein proteostasis axis, showing that the same gene shapes not only social behaviour but also the molecular programs governing stress resistance and lifespan. Because the molecular basis of behaviour is often conserved across taxa and because neuroligins in particular retain conserved structure and function from invertebrates to vertebrate, our findings are positioned to inform future work on the genetics of social behaviour in other organisms, both vertebrate and invertebrate.

## Acknowledgments

This study would not been possible without the continued funding received by the Biotron Institute for Experimental Climate Change Research, a Faculty of Science Core Research Facility at the University of Western Ontario, including the Insect Suite within the Biotron, for housing our flies in climate-controlled walk-in rooms. We are immensely grateful to the essential Flybase [73] (NHGRI awards U$!HG00739 and U24HG010859).

## Funding

This project This project was funded by Internal funding from the University of Western Ontario, and Natural Sciences and Engineering Research Council of Canada Discovery grant 05054-2022 to A.F.S.

## CRediT author statement

Conceptualization, Methodology, Validation, Investigation, Formal Analysis: JMK and RA; Resources, Supervision, Project administration, Funding acquisition: AFS; Software: RA; Writing - Original Draft: JMK; Visualization, Writing - Review & Editing: JMK, RA and AFS.

## Notes

### Competing Interest Statement

The authors have declared no competing interest.

### Summary of Updates

A error was made in Figure 3A - the wrong graphs were shown. This revision now has the representative graphs described in the figure legend and in the text. No change to any other section or text has been done.

